# labapi: a Python object model for the LabArchives electronic lab notebook

**DOI:** 10.64898/2026.09.02.748406

**Authors:** Christoph Li, Josh Lawrimore, Dustin Moraczewski, Adam G. Thomas

## Abstract

`labapi` is a Python library that enables computational workflows to connect to LabArchives’ electronic lab notebook (ELN). Without an Application Programming Interface (API) connection, researchers must manually add workflow outputs through the LabArchives web interface, navigating to the appropriate page and uploading each output so that it appears with the experimental notes that provide context. `labapi` translates the flat LabArchives API into a Python object model following the existing hierarchy of the web interface, allowing workflows to navigate and modify notebook content through familiar paths.

`labapi` enables researchers to build interconnected workflows that both read in and write data to LabArchives’ ELN automatically. Researchers can inspect those outputs in the notebook, and later analysis code can read them back for another stage of analysis.

## Statement of need

NIH intramural policy requires new research to be documented in an approved electronic notebook, and LabArchives is one approved option (NIH Office of Intramural Research, 2024). LabArchives is also used by academic and other research organizations outside NIH (LabArchives, LLC, 2026b). Once uploaded to LabArchives, an output becomes part of the laboratory record beside the notes and revision history that document the experiment (LabArchives, LLC, 2026b).

When a pipeline produces an output to be recorded after every run, using the web interface requires the researcher to return to the appropriate page each time. Researchers can automate that transfer through the LabArchives web API, but doing so requires custom integration code. Researchers navigate the web interface by notebook path; API calls require an internal identifier for each page. Code using the API directly must resolve the path to that identifier and construct the requests that read or write the page (LabArchives, LLC, 2026a). `labapi` performs this work so that researchers can configure workflows with notebook paths. The authors develop two applications built on `labapi` that other NIMH intramural laboratories use in their research: muronto records neuro-behavioral experiment outputs, while `save-my-jupyter` deposits Jupyter notebook snapshots (Lawrimore, 2026; Li, 2026). The intended users are computational researchers and research software developers working in or with experimental laboratories that use LabArchives.

## Software design

`labapi` was designed to simplify working with the LabArchives API and make it familiar to users of the web interface. To provide this familiarity, `labapi` represents the notebook hierarchy shown in the web interface as Python objects. Moving through this hierarchy is essential to working with a notebook, but difficult through the LabArchives API. Helpers such as `dir()` and `page()` ensure that directories and pages exist at a path, retrieving matching nodes or creating missing nodes and parent directories, while `traverse()` resolves only existing paths (Figure 1).

**Figure 1:**
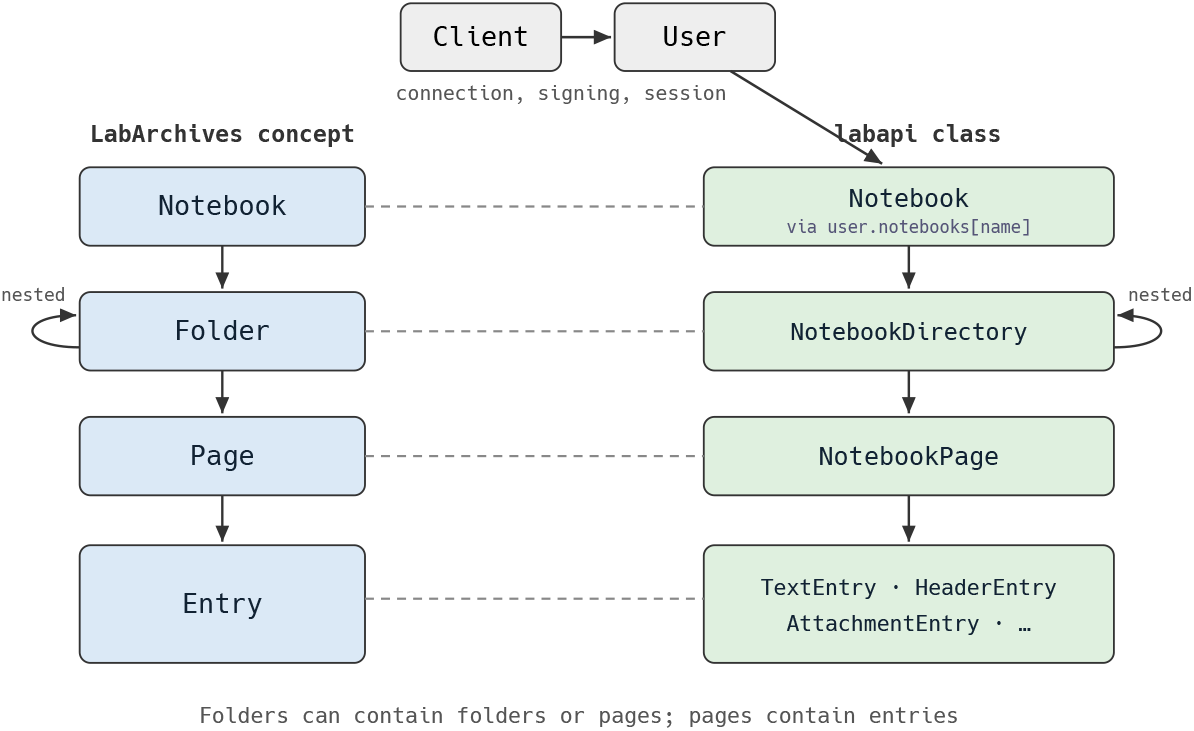
The LabArchives notebook hierarchy and its corresponding `labapi` objects.

Additionally, the design follows the maxim “Parse, don’t validate” (King, 2019). `labapi` parses escaped notebook-path strings into canonical NotebookPath objects that preserve segment and absolute/relative semantics for composition, resolution, containment, and tree traversal.

`create_json_entry()` stores JSON as an attachment for reuse by code and adds a formatted preview for review in the web interface. A library-level meta-entry was considered, but it would break the direct correspondence between `labapi` objects and native LabArchives entries. Preserving the grouping for other tools would then require additional bookkeeping entries, so richer metadata schemes were left to higher-level applications.

Applications built on `labapi` may also need to serve more than one researcher. API credentials are therefore held in a `Client`, separate from each `User` session, so a centralized application can retain multiple sessions without distributing the credentials.

## State of the field

Benchling, RSpace, and eLabFTW pair their APIs with official client libraries (Benchling, 2026; eLabFTW, 2026; Research Space, 2026). LabArchives publishes its API without a public official client library. The standalone LabArchives alternatives in the table do not combine path navigation, entry handling, and attachment transfer in a single client (LabArchives, LLC, 2026a):

**Table 1:** Standalone public LabArchives clients, checked 20 August 2026.

| Project | What it provides | Navigable notebook objects |
| --- | --- | --- |
| `labapi` | General Python client | Yes |
| `labarchives-py` (Cmero, 2022) | Signed generic GET-call wrapper | No |
| `labarchives-js` (Fuschi, 2019) | Login, image-entry search, and attachment-URL helpers | No |
| `labarchives-client` (alaninmcr, 2015) | Unmaintained: no released functionality | No |

Among the standalone projects in the table, `labarchives-py` comes closest. Its single-module implementation signs arbitrary GET requests and returns raw responses, leaving researchers to choose endpoints, parse responses, and implement notebook navigation, entry handling, and file-transfer helpers. Extending it would still have required building the notebook model and higher-level record operations, so `labapi` was developed as a separate client. Specialized ReDBox and MCP integrations also exist, but neither provides a general-purpose, path-oriented Python notebook object interface (Brudner, 2026; University of Technology Sydney eResearch, 2024).

## Example workflow

In the example, a researcher adds `labapi` to a two-stage quality-control workflow using five subjects from the public OpenNeuro dataset `ds000228` (Richardson et al., 2019). The schematic listings use subjects as a placeholder for the five subject identifiers and `compute_qc()` and `summarize()` as placeholders for scientific calculations, so only the high-level LabArchives interaction is shown. Complete setup and entry-handling examples are available in the documentation for working with paths, JSON entries, and example applications. During the first stage, the workflow writes a group label and mean derivative of root mean square variance over voxels (DVARS) for each subject to LabArchives. Mean DVARS is a quality-control value that summarizes how much a functional magnetic resonance imaging (fMRI) scan changes between successive time points (Power et al., 2012). The second stage reads the stored values from LabArchives and produces a cohort summary.

The first stage uses an existing notebook named `Lab QC`, creates or reuses subject pages under `Partly Cloudy QC`, and writes each subject’s data as a JSON attachment with a formatted preview.

~~~
user = labapi.Client().default_authenticate()
qc = user.notebooks[“Lab QC”].dir(“Partly Cloudy QC”)
for subject in subjects:
    qc.page(f”sub-{subject}”).entries.create_json_entry(compute_qc(subject))
~~~

During the second stage, the workflow opens the existing `Partly Cloudy QC` folder with traverse(). Each subject page contains the JSON attachment followed by its formatted preview, so the workflow loads the first entry from every page into `records`.

After loading the five JSON files, `summarize()` produces a cohort summary and figure. The workflow writes both to the `Dashboards/Cohort QC` page.

~~~
notebook = user.notebooks[“Lab QC”]
qc = notebook.traverse(“Partly Cloudy QC”)
records = [json.load(page.entries[0].content) for page in qc.children]
summary, figure_path = summarize(records)
dashboard = notebook.page(“Dashboards/Cohort QC”)
dashboard.entries.create_json_entry(summary)
~~~

Finally, the workflow attaches the figure to the same page.

~~~
attachment = labapi.Attachment.from_file(figure_path)
dashboard.entries.create(labapi.AttachmentEntry, attachment)
~~~

The resulting LabArchives page is shown in Figure 2.

**Figure 2:**
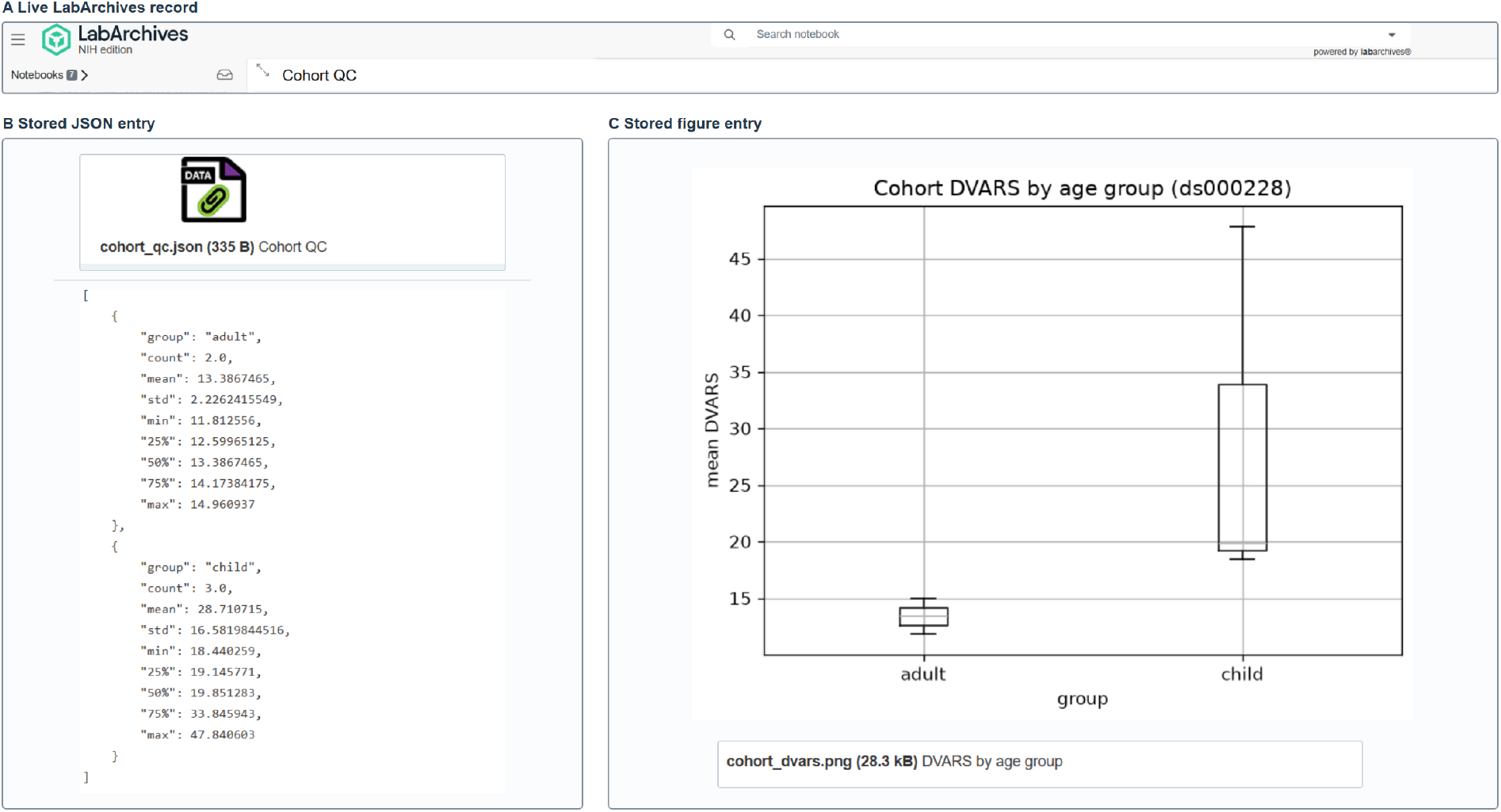
The live LabArchives page containing the cohort summary, a formatted preview of the summary, and the uploaded figure.

## Research impact statement

`labapi` underpins laboratory record-keeping tools in use beyond the authors’ team, including the muronto and save-my-jupyter applications described above (Lawrimore, 2026; Li, 2026). The library’s notebook-backup support was developed in coordination with the Systems Neuroscience Imaging Resource (SNIR), a core facility providing imaging and image-analysis support to NIMH intramural investigators, which uses `labapi` to automate backups of its LabArchives notebooks (nimh-dsst/labapi contributors, 2026). NIH policy also requires intramural researchers to use an approved electronic notebook for new research (NIH Office of Intramural Research, 2024), and LabArchives publishes no official client library; `labapi` fills that gap for laboratories that need to automate this record-keeping.

## Availability

The source and documentation are available at https://github.com/nimh-dsst/labapi and https://nimh-dsst.github.io/labapi/v1.2.0/, respectively. Version 1.2.0 is archived on Zenodo (Li et al., 2026) and released under the MIT License. `labapi` is also published on PyPI, can be installed with `pip install labapi`, and requires Python 3.10 or later.

The repository includes end-to-end examples, credential-free unit tests, and opt-in live integration tests.

## Author contributions

Author contributions are described using the CRediT taxonomy. Christoph Li led software development (Software) and prepared the original manuscript draft (Writing – original draft). Josh Lawrimore conceived the project (Conceptualization) and contributed code and code review (Software). Dustin Moraczewski and Adam Thomas provided project direction and oversight (Supervision). Josh Lawrimore, Dustin Moraczewski, and Adam Thomas reviewed and edited the manuscript (Writing – review & editing).

## AI usage disclosure

### Nature and Scope

The authors designed `labapi`‘s foundational architecture and implemented its core. Generative AI subsequently assisted with development, testing, maintenance, release work, project documentation, and manuscript preparation. It generated code, contributed to tests, opened pull requests and issues, and drafted and edited documentation and manuscript text.

### Author Review

The authors retain final control over the software’s scope, behavior, and public interface and take responsibility for the software and manuscript. They reviewed and accepted AI-assisted software and documentation changes through public pull requests and reviewed and revised all AI-assisted manuscript text. GitHub Actions runs unit tests on every push and pull request and live LabArchives integration tests when triggered.

### Tools

Anthropic’s Claude (Opus 4.6, Opus 4.8, Opus 5, Sonnet 4.6, Sonnet 5, and Fable 5) and OpenAI’s Codex (GPT-5.4, GPT-5.5, and GPT-5.6 Luna, Sol, and Terra), the latter accessed through the HHS Enterprise agreement.

## Acknowledgements

This research was supported in part by the Intramural Research Program of the National Institutes of Health (NIH). The contributions of the NIH author(s) were made as part of their official duties as NIH federal employees, are in compliance with agency policy requirements, and are considered Works of the United States Government. However, the findings and conclusions presented in this paper are those of the author(s) and do not necessarily reflect the views of the NIH or the U.S. Department of Health and Human Services (HHS). NIH support was provided under NIMH project ZICMH002960. This project has been funded in whole or in part with federal funds from the National Cancer Institute, National Institutes of Health, under Contract No. 75N91019D00024. The role of these funding sources was limited to supporting the researchers through the NIH Intramural Research Program and the listed NCI contract. The content of this publication does not necessarily reflect the views or policies of the Department of Health and Human Services, nor does mention of trade names, commercial products, or organizations imply endorsement by the U.S. Government.

